# Coral restoration is only practical when local human pressure levels are low

**DOI:** 10.1101/2024.11.26.622858

**Authors:** Cole B. Brookson, Ariel Greiner

**Affiliations:** Department of Epidemiology of Microbial Diseases, Yale University; Department of Biological Sciences, University of Alberta; Department of Biology, University of Oxford, Oxford, UK; Center for Infectious Disease Dynamics and Department of Biology, Pennsylvania State University

**Keywords:** Coral Reef, Restoration, Mathematical Modelling, Nutrient Enrichment, Fishing, Management Planning

## Abstract

Coral reefs are threatened by interacting local (fishing, coastal development) and global (climate change) stressors that degrade coral cover. Adding coral to the reef (coral restoration) is gaining prominence as a management strategy, however it is unclear whether restoration alone can ensure self-sustaining high coral cover under interacting stressors. We assess how much restoration is needed for a self-sustaining healthy coral reef using a mathematical model and real-world restoration case studies. We find that a reef with low macroalgal cover and high coral recruitment requires low levels of restoration to ensure self-sustaining high coral cover but as nutrient enrichment and fishing increase, more restoration is required. Thus, if the reef is facing high local human stressors, restoration is unlikely to ensure self-sustaining high coral cover, suggesting that under such conditions conservation resources should first be allocated towards reducing the impact of stressors, prior to any restoration.

## 1 Introduction

Coral reefs face numerous imminent threats to their survival, both at the global level from bleaching events, ocean acidification, and damage from severe storms [1, 2] as well as the local level from overfishing [3], nutrient enrichment [4], and sedimentation [5]. These stressors have halved living coral levels worldwide since the mid-20th century, and could lead to between 70–99% of reefs experiencing bleaching-related degradation by 2100 [6, 7, 8]. Efforts to combat declines in coral health include reducing fishing via gear restrictions or periodic limits, reducing erosion through runoff control, and coral restoration [9], where coral fragments, coral larvae or structures/substrate are artificially added to a reef in an effort to directly increase coral cover [10].

Attempts to assist coral in its regrowth are hindered by the fact that the same stressors which suppress coral often allow benthic organisms such as macroalgae, which directly compete with coral, to grow more easily. The increased ability of these organisms to compete with coral under these stressors can cause reefs to shift from coral-dominated to non-coral dominated ecosystems [11]. This is particularly concerning because coral reefs are thought to have multiple “stable states”, including macroalgal-dominated states [12, 13, 14] which support low fish biodiversity [15].

Multiple stable states (MSS) exist when a single ecosystem can support two or more stable communities [12]. In coral reef ecosystems, the two main stable states discussed are coral-dominated states and macroalgal-dominated states [16, 12, 13]. Coral reef MSS were first modeled by [13], but further theoretical research has extended this work to understand how processes such as larval dispersal [17, 18, 19] and other stressors such as sedimentation and nutrient enrichment [14] affect the stability of the coral-dominated state. Some have questioned the presence of MSS on reefs, arguing that coral-dominated and macroalgal-dominated states exist along a stable continuum instead [20, 21], but empirical data from the Indo-Pacific and the Caribbean [22, 23, 24] and experimental evidence from the Indo-Pacific [25] support predictions consistent with the existence of MSS on coral reefs. Coral reef MSS dynamics are important for management planning, as they imply the existence of thresholds that should be considered when designing management plans [26].

MSS are particularly relevant in the context of coral loss and restoration because coral competition with algae for space is a known threat to the growth and settlement of newly dispersed coral [27]. Widespread restoration activities have attempted to reduce coral loss [9, 10] but they are incredibly costly [28] and have shown mixed success [29, 10]. Large-scale restoration studies have not been feasible due to the difficulty of adding more than a small percentage of coral cover to a large reef system [10]. Additionally, even in restoration studies that did find high survivorship of out-planted coral, bleaching events remain a challenge [30], and maintenance (e.g. clearing macroalgae from nursery corals [31]) or removing corallivores [32] was required.

These shortcomings highlight a key factor for managers: the importance of theoretical work *prior* to restoration activities to assess the viability of the plan [10]. Coral reef restoration may be an appropriate conservation management plan for certain reefs but not all [33]. Coral reef managers should use feasibility modeling to determine when restoration is appropriate, as in other conservation initiatives. Well-documented examples of this type of “theory-before-practice” approach include population viability analyses [e.g., 34, 35, 36] which have contributed to the famously successful re-introduction of species like wolves and Peregrine falcons [37] and management strategy evaluation methods in fisheries [e.g., 38, 39, 40]. In the case of coral reef conservation, we argue here that we can use simplified models of coral reef dynamics to take the internal dynamics and stressor levels of a coral reef into account so as to assess which reefs may be most benefited by restoration methods. We use a simple model of coral reef cover to provide broadly applicable guidance for successful coral restoration. We assess when restoration may be feasible by simulating how coral cover responds, over a long time period, to different combinations of stressor levels. This helps assess whether achieving and maintaining a coral-dominated stable state may be possible at different levels of coral restoration, or if the system is likely to regress to a degraded state. We modify a standard model from [13] and consider stressors including fishing levels, nutrient enrichment, and health of connected reefs. We then use two case studies as examples for how this modeling approach could be applied in specific situations.

Overall, our goal is to provide insights and guidelines for applying models to better inform the rapidly growing field of coral reef restoration.

## 2 Methods

### 2.1 Overview

To assess which conditions support restoration as a reasonable management method for coral reefs, we developed a simple model to determine the minimum coral cover that restora-tion efforts would need to achieve, under varying levels of interacting stressors and coral larval recruitment, to allow a reef to trend toward a coral-dominated state with coral cover *C >* 30% and macroalgal cover *M <* 30% (i.e. a self-sustaining healthy coral reef state). In this study, we focus on coral restoration efforts that directly add coral fragments or coral larvae to the reef, not considering those that only add structures or alter substrate to facilitate the growth or settlement of additional coral [10]. We use a linear stability analysis to find stable equilibria (attractors) of the system. Then, we simulate a range of trajectories from different initial conditions and parameter combinations (representing different stressor levels) to determine the basins of attraction associated with each attractor. These basins of attraction allow us to understand under which initial conditions and parameter combinations the system approaches a coral-dominated attractor, and thereby could attain and maintain a coral-dominated state (i.e. restoration “success”). Finally, we used real-world restoration examples from the literature as case studies to demonstrate the application of this approach in real systems, and make quantitative predictions of restoration success.

### 2.2 Model

Our mathematical model of a reef is based on the system of ordinary differential equations in Mumby et al. [13], to which we added an explicit coral larval recruitment term to allow us to model larvae dispersing and settling into the focal reef from connected reefs (based off of the Elmirst et al. [17] model). The model describes how the coral cover (*C*), macroalgal cover (*M*) and turf algae cover (*T*) change over time in a single reef. The model takes the form of:

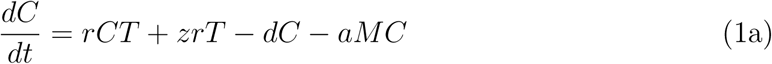

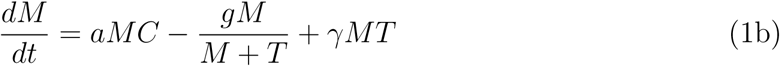

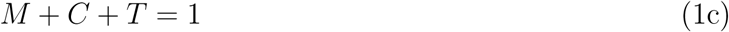

Equation (1a) describes the rate of change in coral cover, *C* and Equation (1b) describes the rate of change in macroalgal cover, *M* . The model describes how herbivorous grazing on macroalgae and turf, natural coral mortality, coral larval recruitment, and spatial competition processes on the reef change the relative cover of coral (*C*), macroalgae (*M*), and turf algae (*T*) through time (see Figure 1). The amount of *C* on the focal reef increases as mature coral and coral larvae from the focal reef out-compete existing turf algae at rate *r* and as coral larvae recruit onto the reef from connected reefs at rate *z*. *z* is higher when more coral larvae are recruiting into the focal reef, indicating either healthier connected reefs or more connected reefs (due to changing ocean current patterns). We use a simple, non-time varying functional form for coral recruitment (*zrT*) following [17, 19] and because [41] found that approximating time-varying dispersal with constant average dispersal rates was appropriate (as the average rate of external recruitment was the most important factor for determining the dynamics of the system). Coral can be removed from the system either through a natural mortality rate *d* (*d* = 0.24), or they can be out-competed by macroalgae, at rate *a*. If the reef experiences high levels of nutrient enrichment, macroalgae’s competitive advantage over coral grows via an increase in *a* [4]. Since nutrient enrichment has been found to increase the growth rate of macroalgae and turf algae [42, 43, 14, 44] and decrease the growth rate of coral, we decided to model increasing nutrient enrichment as increasing *a* and not *γ*. Mature macroalgae and macroalgae gametes out-compete turf algae at a rate *γ* (*γ* = 0.77). Macroalgae and turf are removed from the reef by grazing processes at the rate 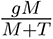. We assume that grazing on the reef can be from any herbivorous species (e.g. parrotfish) that can reasonably be assumed to graze indiscriminately on both turf and macroalgae. Grazing makes room for dispersing coral larvae to land and settle (since, as described in Equation Equation (1c), there is no “empty” space in this model). The grazing rate *g* on a reef is dependent on which herbivorous taxa are present, in what abundance and on their consumption rate of different benthos. As we are not focused on any particular reef system, we assume indiscriminate grazing on macroalgae and turf algae following other similar models [13, 17, 19]. We included recruitment of coral larvae since that is a major factor driving the success of coral [17, 19], but here assume that the only macroalgae gametes recruiting onto this reef are from the focal reef (only able to disperse short distances) [45, 46].

**Figure 1:**
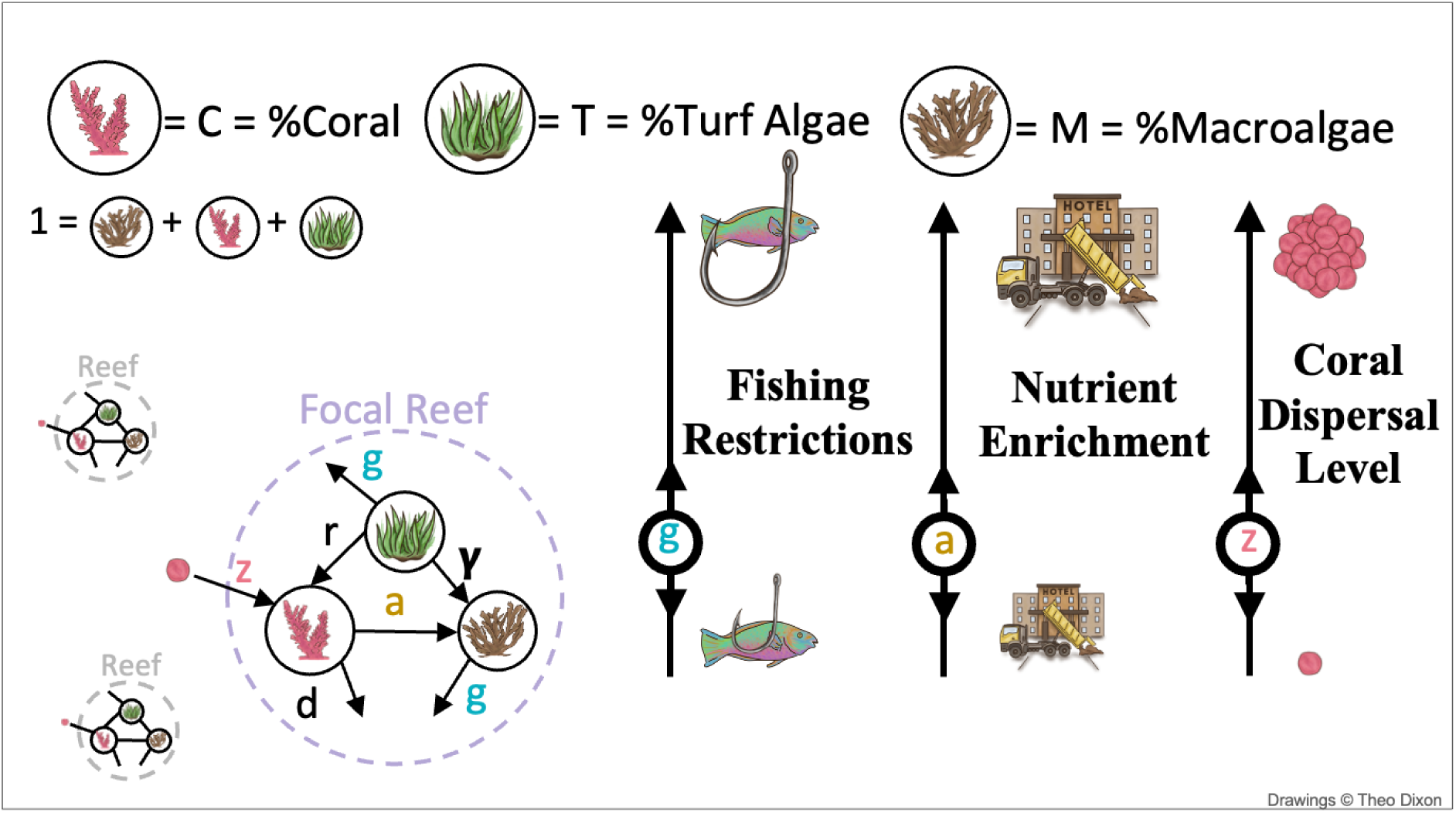
Conceptual figure of the model and how it represents the coral reef. We consider the reef as being covered entirely by coral (C), macroalgae (M) and turf algae (T). For simplicity, we say that (T) represents turf but this state variable essentially represents any space-filling benthic organism that can be settled on by coral larvae and consumed by herbivores (in some similar models it represents free space - see [68]). The focal reef is a discrete area that may be surrounded by other discrete reefs from which coral can recruit from. The transition between the three types of cover is modulated by the lettered parameters next to the arrows. The process of coral growth over turf algae (*r*), macroalgal growth over coral (*a*), and recruitment of coral larvae to the focal reef (*z*) and natural coral mortality (*d*) describes coral growth, while macroalgae can outgrow turf algae (*γ*). The colours of the different anthropogenic effects relate to the parameters in the model coloured the matching colour. For more detail see SI methods.

We wanted to assess how different stressors would change restoration success, so we simulated changes in the rate of macroalgal-coral competition *a*, coral larvae recruitment *z* and grazing rate *g* (Table 1) to approximate changes in nutrient enrichment levels, health of connected reefs, and fishing levels (respectively) (Figure 1). We varied these parameters continuously, but for some visualizations they are binned as ‘low’, ‘medium’, and ‘high’, and performed a fully factorial comparison across all three variables. We also wanted to assess if the percentage cover of *C*, *M*, and *T* affected restoration success due to inherent non-linearities, so we compared a thorough sampling of possible starting combinations of the three state variables and observed how that impacted the final state of the reef. From this, we determined, at each parameter combination, what the minimum initial coral cover value was that would lead the focal reef to trend to a self-sustaining healthy coral reef state from any starting *C* cover value (Figure 2). We then quantified how that minimum initial coral cover value changed if we focused only on low, medium or high initial macroalgal cover values (Figure 3).

**Figure 2:**
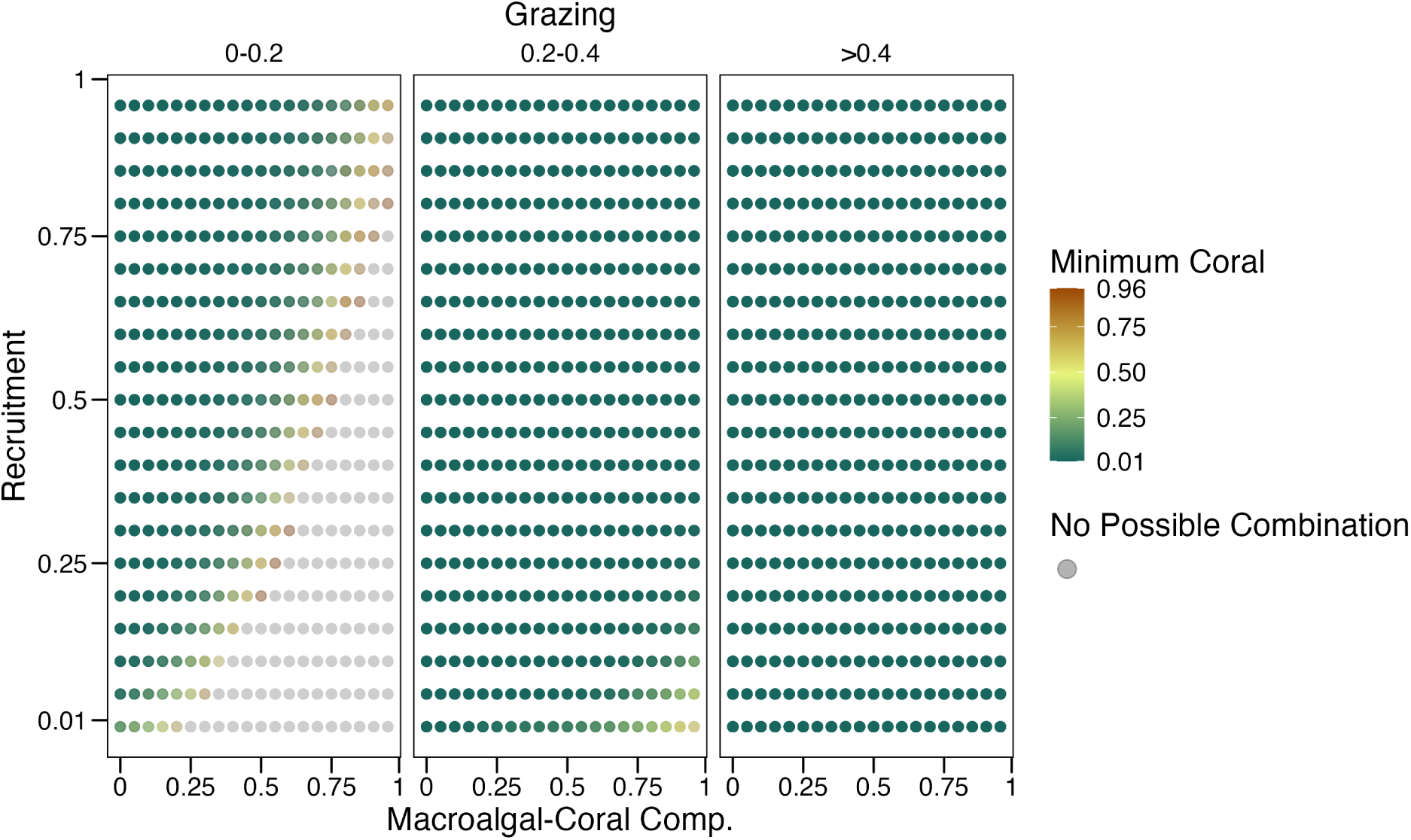
Each point represents the minimum amount of initial coral cover for the given parameter combinations, such that all initial values of macroalgae result in the reef being in a “healthy self-sustaining coral reef state” (i.e., trending towards a stable state with *>*30% coral cover & *<*30% macroalgae cover). Light grey points indicate that no possible parameter combination or initial condition results in the reef being in a “healthy self-sustaining coral reef state”.

**Figure 3:**
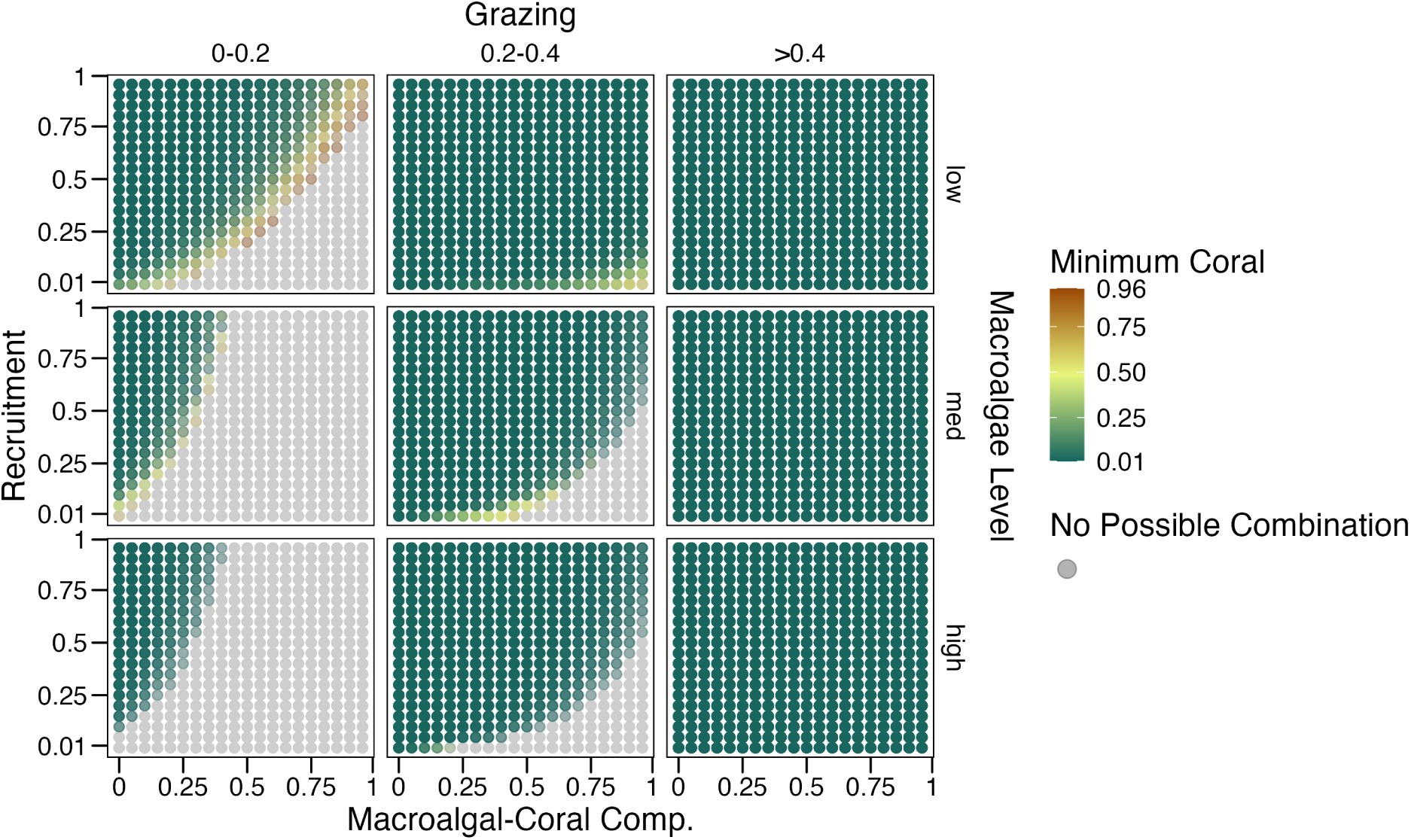
Each point represents the minimum amount of initial coral cover for the given parameter combinations, such that certain initial values of macroalgae result in the reef being in a “healthy self-sustaining coral reef state” (i.e., trending towards a stable state with *>*30% coral cover & *<*30% macroalgae cover). The values on the right y-axis depict whether the initial values of macroalgae considered are low (0-33%), medium (33-66%) or high (>66%). Light grey points indicate that no possible parameter combination or initial condition results in the reef being in a “healthy self-sustaining coral reef state”.

**Table 1:** Parameter values and details for the set of differential equations used to simulate the coral reef system. Variables that are considered levers for conservation action (*z*, *a*, and *g*) are explained with respect to their usefulness in conservation.

| Model Parameters |  |  |  |
| --- | --- | --- | --- |
| Term | Description | Values | Restoration Importance |
| $r$ | Rate of coral larval overgrowth on turf algae | 0.55 | |
| $z$ | Recruitment rate of coral into the reef | 0-0.99 | The health of nearby reefs will strongly affect the success of coral recruitment into the focal reef, and hence the amount of new coral to supplement the recently transplanted coral. |
| $d$ | Natural mortality rate of coral | 0.24 | |
| $a$ | Rate of macroalgal-coral competition | 0-0.99 | The rate at which macroalgae will succeed in overgrowing coral changes with increases in nutrient influx to a reef [17]. If a reef has high nutrient enrichment levels, macroalgae will overgrow coral at higher rates. |
| $g$ | Rate of grazing on algae | 0-0.99 | Herbivore numbers on reefs will impact restoration since if grazing is low or absent, there is no removal of macroalgae. [17]. |
| $\gamma$ | Rate of macroalgae overgrowth on turf algae | 0-0.77 | |

### 2.3 Case Studies

We used two restoration case studies as examples for how this model could prove useful in a real-world context. To do this, we used the model (Equations (1a–1c), Figure 1) to simulate reefs based on empirically-derived parameters for the case studies to determine whether restoration activities changed reefs’ tendencies to attain a self-sustaining healthy coral reef state. We used data from [47], who tracked not only the restoration activities of sites, but also the general conditions of the region containing the sites. We compared two different case studies: the New Heaven Reef Conservation Program (NHRCP) on the Thai island of Koh Tao, and The Nature Conservancy project (TNC) on St Croix, US Virgin Islands. We compiled data from each of these programs and translated the information about the reefs into ranges of parameter values that could be used in our model. For coral cover and macroalgal cover, we gathered proportions directly from the project data [47], but for other important parameters in the model, we inferred from categorical information on each study [48, 49]. We used information on herbivore abundance or fishing level to inform our grazing parameter, *g*, and immigration level of coral larvae to inform our coral larval recruitment measurement, *z*, (Table 2). We used information about the state of the region itself from [47] to parameterize the macroalgal-coral competition parameter, *a* (e.g. for NHRCP, [47] stated that the reefs were under stress from terrestrial run-off). Once we extracted the parameter values, we simulated each of the case studies across a factorial set of simulations across possible parameter combinations. This allowed us to assess the new starting point for the system and determine whether the increase in coral cover attained through the restoration program described, without altering any of the stressors in the region, would allow for the system to attain a self-sustaining healthy coral reef state in the future.

**Table 2:**
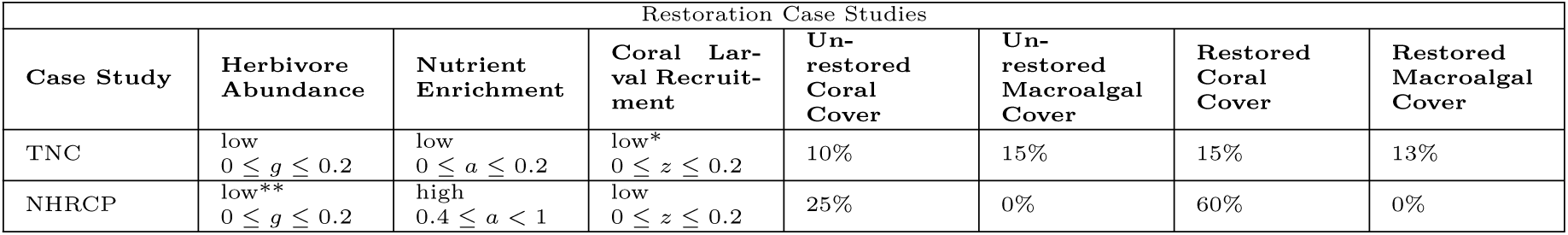
Information from our case studies, translated into model parameters. *Too low to measure pre- and post-restoration. **Assuming low based on inference of high fishing rates from other studies in this area [48, 49].

## 3 Results

We quantified how the minimum initial coral cover required to achieve and maintain a self-sustaining, coral-dominated state (defined as a reef with coral cover *C >* 30% and macroalgae *M <* 30%) varies across grazing, coral larval recruitment, and macroalgal–coral competition, and found that the required minimum is lowest when grazing is high, recruitment is high, and competition is low, whereas under low grazing the threshold often rises above 70% or becomes unattainable in the explored parameter space (Figure 2; cf. [50]).

Initial macroalgal cover substantially modifies these thresholds. Low initial macroalgal cover reduces the initial coral cover needed to reach the healthy state, while higher initial macroalgal cover increases the required initial coral cover and, in some regimes, eliminates the possibility of attaining a healthy endpoint altogether (Figure 3). Across parameter sweeps, varying macroalgal–coral competition and larval recruitment shifts the attainable region smoothly for both the minimum grazing/recruitment values and the requisite initial coral and macroalgal cover levels (Figures S1 to S4). In contrast, varying the grazing level leads to pronounced threshold-like differences in the coral cover required to reach a healthy endpoint (apparent as sharp transitions across grazing levels in Figures 2 and 3).

Applied to two case studies (TNC, NHRCP; Table 2) using their reported final (“restored”) covers and parameterizations, the model predicts a low likelihood of achieving and maintaining a healthy state from 15% (TNC) and 60% (NHRCP) restored coral cover under the specified grazing, competition and recruitment conditions (Figure 4C&D). We also observe that the addition of coral in both systems is unlikely to have changed the overall, long-term outcome of the system (comparing before and after), even in the case when it substantially increased the coral cover, unless it was accompanied by stressor management (Figure 4)

**Figure 4:**
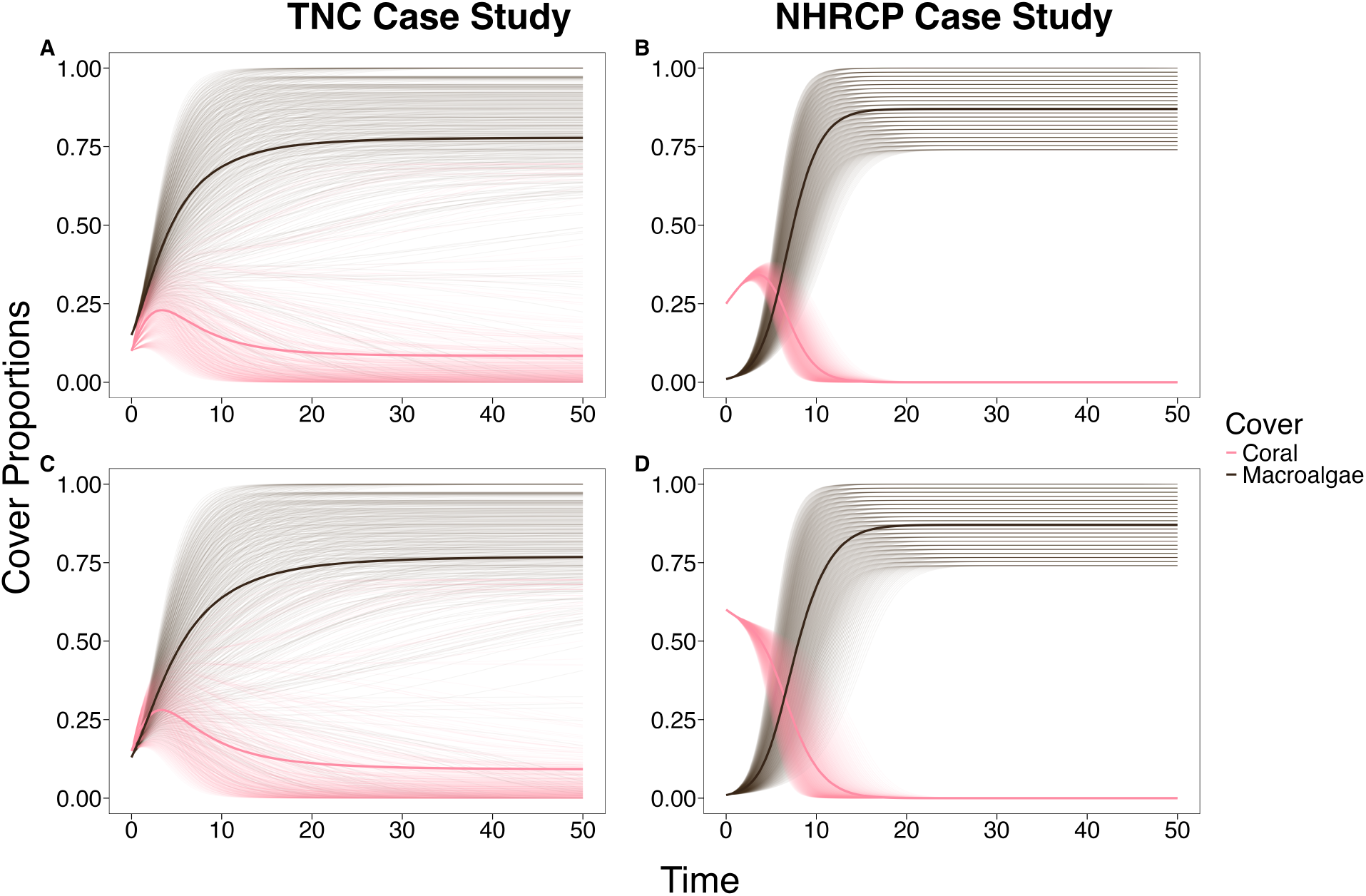
Restoration case studies before coral restoration (first row) and after (second row). A) and C) depict the the St. Croix reefs in the Nature Conservancy (TNC) case study, and B) and D) depict the New Heaven Reef Convservation Program (NHRCP) on the Thai island of Koh Tao.

## 4 Discussion

We found that the minimum amount of initial coral cover needed to achieve and maintain a self-sustaining healthy coral reef (i.e., trending toward a coral-dominated stable state with *C >* 30% and *M <* 30%) is lowest when the grazing rate is high, coral larval recruitment rate is high and macroalgal-coral competition is low (Figure 2). At low grazing rates, as the macroalgal-coral competition value increases and the coral larval recruitment rate decreases, the minimum initial coral cover value needed for the reef to be able to achieve and maintain a healthy state may increase to *>* 70% (which is unrealistic in most real systems [50]) or there may be no sufficient minimum initial coral cover level (Figure 2). These results are echoed in the few small-scale restoration studies which comment on the environmental condition and the amount of other benthic organisms on the reefs throughout the restoration process, which find that reefs with high immigration and connectivity to thriving reefs as well as reefs with low nutrient enrichment or fishing pressures can benefit from restoration [51, 52, 47].

Even at reasonably high levels of coral cover (15% and 60% respectively), neither of the two case studies (TNC, NHRCP; see Table 2) appears likely to achieve and maintain a coral-dominated state in the context of this model because of low grazing and low coral larval recruitment (Figure 4). Given these parameter values and macroalgal cover in the TNC case study, restoration efforts in this region would need to increase the coral cover to 30-40% for the reefs to trend to a healthy state and 50-60% if the macroalgal cover were to increase (Figures 2 and 3). For NHRCP, without changing the parameter values or final macroalgal cover, increasing the coral cover of the system would never allow the system to achieve and maintain a healthy state; managers would need to first reduce the nutrient enrichment levels or find some way to increase the coral larval recruitment (e.g. through increasing the health of coral reefs that send coral larvae to these reefs) (Figure 3).These results indicate that both regions should pursue complementary management alongside coral restoration (e.g., wastewater management to reduce enrichment, fisheries management to increase key herbivores, actions to improve coral cover on connected source reefs after identifying larval sources). However, it is worth mentioning that since the NHRCP reefs were described as having very low macroalgal cover before and after restoration, it is possible that the macroalgal cover level would remain low into the future on these reefs. This is contrary to what our model predicts (Figure 4B&D). It would be worth returning to these sites and measuring the present-day benthic cover (and the coral recruitment, herbivore grazing and nutrient enrichment levels) and comparing it with these results to assess whether the predictions from this model manifested in the system. This additional data would allow us to adjust the model (if necessary) to fit these systems better.

These results encourage paying specific attention to grazing levels on a reef because model behaviour changes most abruptly across grazing levels while recruitment and competition exert smoother, more incremental effects (Figures S1 to S4). In particular, grazing rates on algae (and other benthic substrates) should be explicitly measured so that we do not need to rely on fishing level estimates (Table 2) or grazer biomass levels to approximate it [53]. Therefore, our results encourage restoring fish biomass to an acceptable level on a reef prior to any restoration activities, as this may be critical to allow reefs to recover to a healthier state. This does not diminish the value of increasing larval supply or reducing macroalgal–coral competitive advantage, but it suggests that those levers are more likely to pay off when herbivory is first restored to functional levels (Figures S1 to S4).

This model was developed from other models (most notably, [13]) whose conclusions have been shown to fit real-world systems in the Caribbean [11, 22, 54] and in the Indo-Pacific [24, 25, 23, 55]. Because the base model [13] was built to investigate grazing effects, our finding that varying the grazing parameter can strongly alter restoration success (more than any other parameter; compare Figure 2 with Figures S1 and S2) may be accentuated and merits field tests. Nevertheless, multiple empirical studies across regions have independently underscored the central role of herbivory in sustaining coral-dominated states [56, 57]. Thus the qualitative conclusion—restore herbivore function early—remains robust and worth testing via targeted field experiments that manipulate herbivory and enrichment while tracking recruitment at management-relevant scales. Equally, where local nutrient inputs can be feasibly reduced, managers should expect synergistic gains because limiting macroalgal competitive advantage lowers the initial coral cover threshold for recovery in our simulations (Figure 3).

In general, additional data could have improved this study. In particular, we were unable to find long-term data from coral restoration studies in a form that we could use to parameterize this model. [58] provide large amounts of useful data but since it assesses the effectiveness of the addition of structures, not the addition of coral larvae or fragments themselves, it was not within the scope of this study. Most case studies report short-term survival of coral out-plants (coral fragments grown in nurseries that are planted back onto degraded reefs to aid in ecosystem restoration) rather than time series of reef-wide benthic cover, long-term post-restoration coral cover, or environmental conditions before/after restoration. In particular, having benthic cover and environmental data from a coral restoration initiative that found long-term success would have been invaluable. This lack of relevant data is attributable both to the relatively recent proliferation of coral restoration projects, which restricts the duration of available time series, and to the limited collection of model-relevant parameters in empirical field studies. There have been studies advocating for these parameters to be measured in the field as they are relevant for diagnosing reef resilience (e.g. [59] - coral recruitment rate, herbivorous fish grazing rates) and many studies have demonstrated how to measure them (e.g. herbivorous grazing rates - bite rates, herbivore visitation rates [25], but they are more labour intensive and may require repeated dives at the same site within a short period of time. Thus, it would be helpful if there existed more information on how to translate commonly measured values (e.g., herbivore abundance/density, photos of benthic cover over time) into rate parameters (e.g., grazing on macroalgae/turf algae, coral–macroalgal competition, natural coral mortality) relevant to resilience assessments and usable in models like the one used in this study.

This model varies initial coral cover values to simulate possible endpoints of restoration initiatives, implicitly assuming the intervention succeeded in raising coral cover to those levels. If some out-plants fail (e.g., due to restoration technique or existing environmental stressors), the resulting post-failure cover can be treated as the new initial coral cover level and re-evaluated in the same framework. In this way, the results of this study could also be used to assess the effectiveness of management initiatives focused on reducing human-induced coral damage (e.g. stemming from anchoring, blast fishing, direct damage from tourists, etc.). The results that we discuss from this model focus on whether the focal reef will trend towards a healthy coral cover from a particular initial coral cover. However, whether the reef will achieve a healthy coral cover is dependent on the environmental conditions of the reef that define the parameters in the model remaining constant and assumes that no other factors will cause changes to the benthic cover. This is unlikely to be the case (due to constant environmental change and increased frequency of bleaching events and cyclones), however the formulation of this model would allow one to re-assess whether the focal reef is likely to achieve this healthy coral cover at new parameter and initial benthic cover values. We focused on coral cover as the success metric given its importance for reef health [60, 61], but we do recognize that other metrics (e.g., structural complexity, biodiversity, ecosystem service provision) may be equally or more relevant depending on objectives [53, 15, 62, 63, 59]; future work should test whether conclusions change under alternative metrics.

In an era of rapid change, integrating models with site-specific data and co-designing analyses with managers can help prioritize interventions and avoid investments in conditions unlikely to yield self-sustaining coral dominated states. Using models to assess management options in advance is common in other domains (species introductions, fisheries) [64, 65]; similarly, the present framework can help determine when coral restoration alone is sufficient versus when complementary actions should precede or accompany restoration efforts.

## 5 Conclusion

Overall, our results show that the best method for achieving self-sustaining healthy coral reefs may be to first reduce nutrient enrichment on reefs, increase herbivore grazing rates, and increase coral cover on well-connected reefs (through similar methods) to increase coral larval recruitment values [66, 19] (Figures 2 and 3). Pursuing these actions before pursuing coral restoration efforts could help avoid wasting resources on reefs with little to no herbivore abundance or coral larval recruitment–which is paramount for conserving resources while effectively protecting coral reefs. Our results indicate that the best candidates for coral restoration are reefs that have low coral cover but are otherwise unaffected by local human pressures – i.e. reefs that have recently been hit by cyclones or bleaching events but otherwise have low stressor levels. However, even in such an ideal case it is important to consider whether the reef with the newly dead coral remains healthy enough to recover on its own and whether adding more coral would be a waste of effort (e.g. important to use coral that are thought to be resilient/resistant to bleaching in areas thought to be at high risk of future bleaching events, [67]). The process of identifying potential candidate coral reefs for restoration should include developing a mathematical model that incorporates more specific knowledge of a given reef as well as the capacity of the community and other involved stakeholders to maintain proposed management efforts (including restoration).

## Acknowledgement

The authors thank Dr. Alexandra Davis for helpful conversations and edits during the preparation of the manuscript and Theo Dixon for their illustrations for Figure 1. CBB was supported by an NSERC PGS-D fellowhip and partially supported by an NSF Biology Integration Institute grant (NSF DBI 2213854). AG was supported by an NSERC CGS-D fellowship and an NSERC Postdoctoral fellowship.

## 6 Data Availability Statement

Data sharing is not applicable to this article as no new data were created or analyzed in this study. All code to reproduce the analyses here can be found on GitHub: https://github.com/colebrookson/coral-restoration/releases/tag/v1.0.0

---

**Figure S1:**
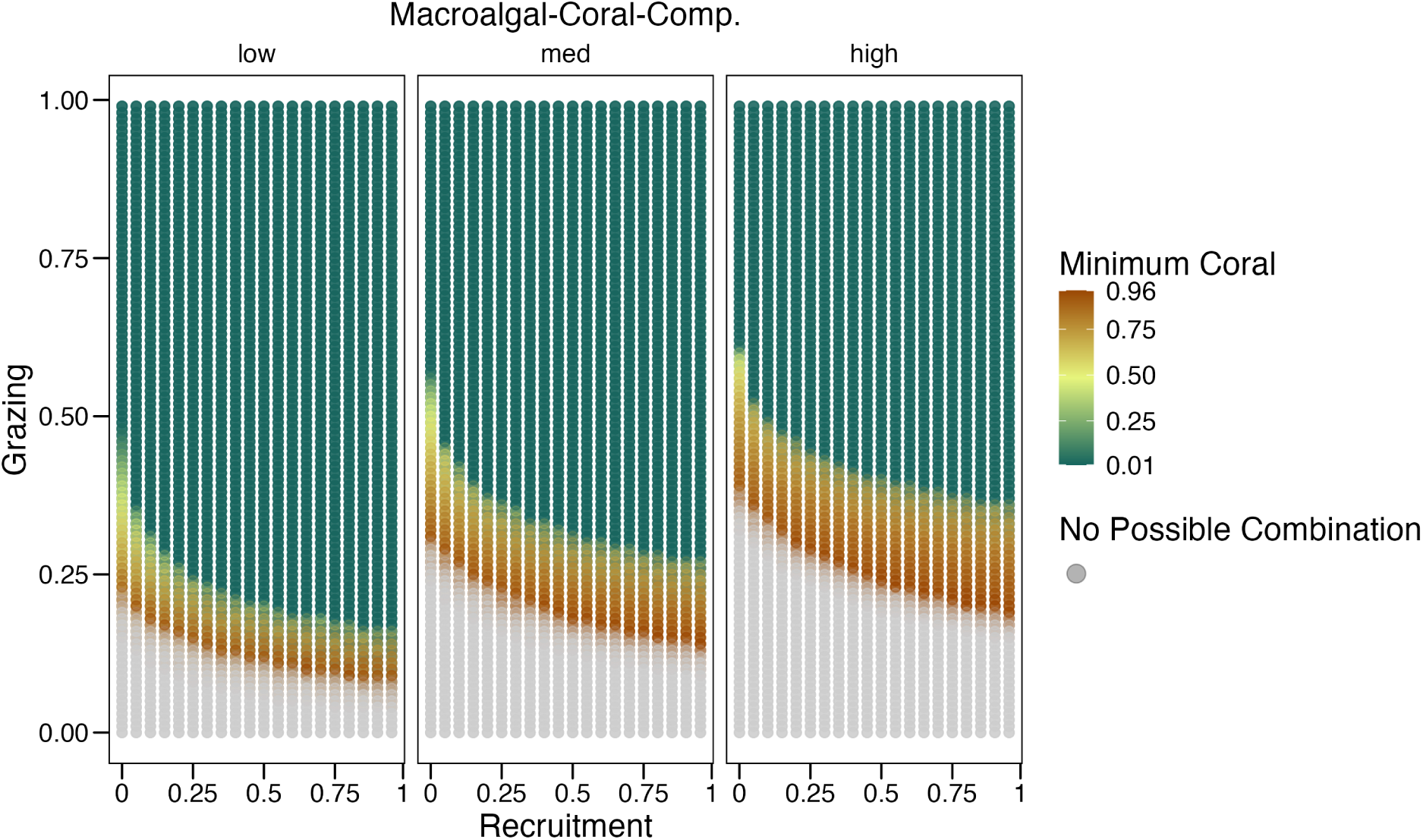
Minimum amount of coral cover (percentage) required, given a particular parameter combination, for a given trajectory to result in the reef being in a “healthy self-sustaining coral reef state” (i.e., *>*30% coral cover & *<*30% macroalgae cover). Light grey points indicate that no possible parameter combination or initial condition will result in the reef being in a “healthy self-sustaining coral reef state”.

**Figure S2:**
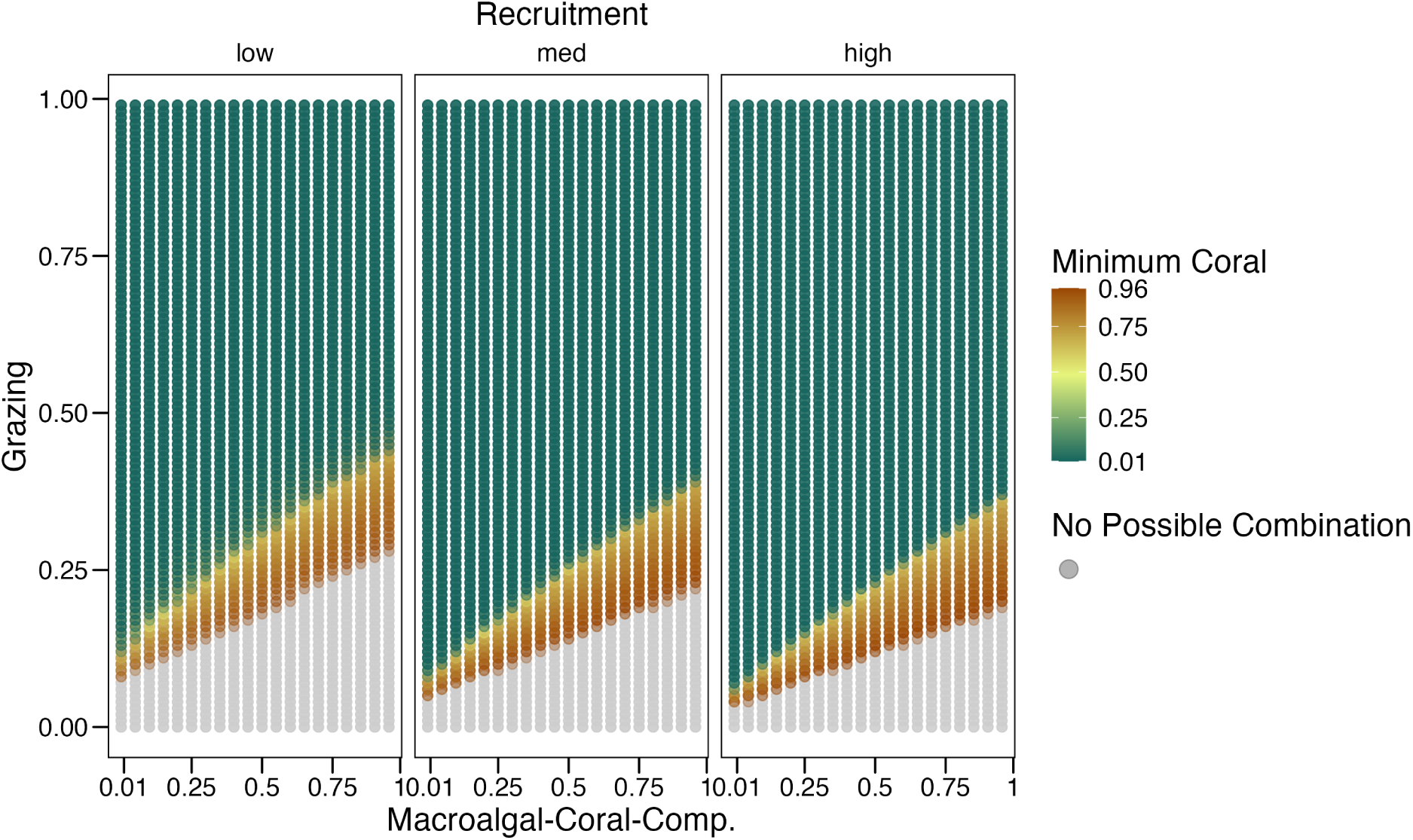
Minimum amount of coral cover (percentage) required, given a particular parameter combination, for a given trajectory to result in the reef being in a “healthy self-sustaining coral reef state” (i.e., *>*30% coral cover & *<*30% macroalgae cover). Light grey points indicate that no possible parameter combination or initial condition will result in the reef being in a “healthy self-sustaining coral reef state”.

**Figure S3:**
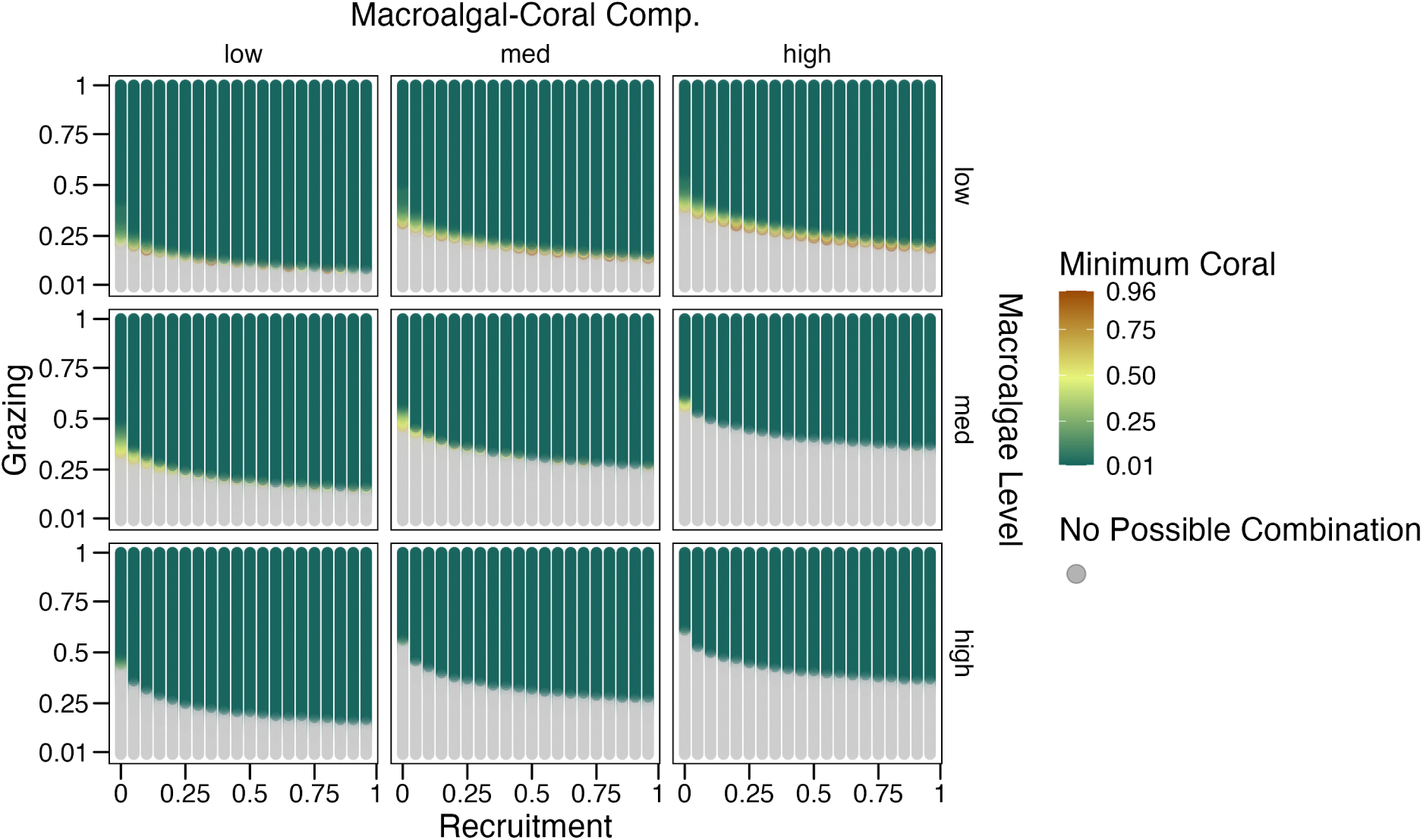
Each point represents the minimum amount of initial coral cover for the given parameter combinations, such that certain initial values of macroalgae result in the reef being in a “healthy self-sustaining coral reef state” (i.e., trending towards a stable state with *>*30% coral cover & *<*30% macroalgae cover). The values on the right y-axis depict whether the initial values of macroalgae considered are low (0-33%), medium (33-66%) or high (>66%). Light grey points indicate that no possible parameter combination or initial condition results in the reef being in a “healthy self-sustaining coral reef state”.

**Figure S4:**
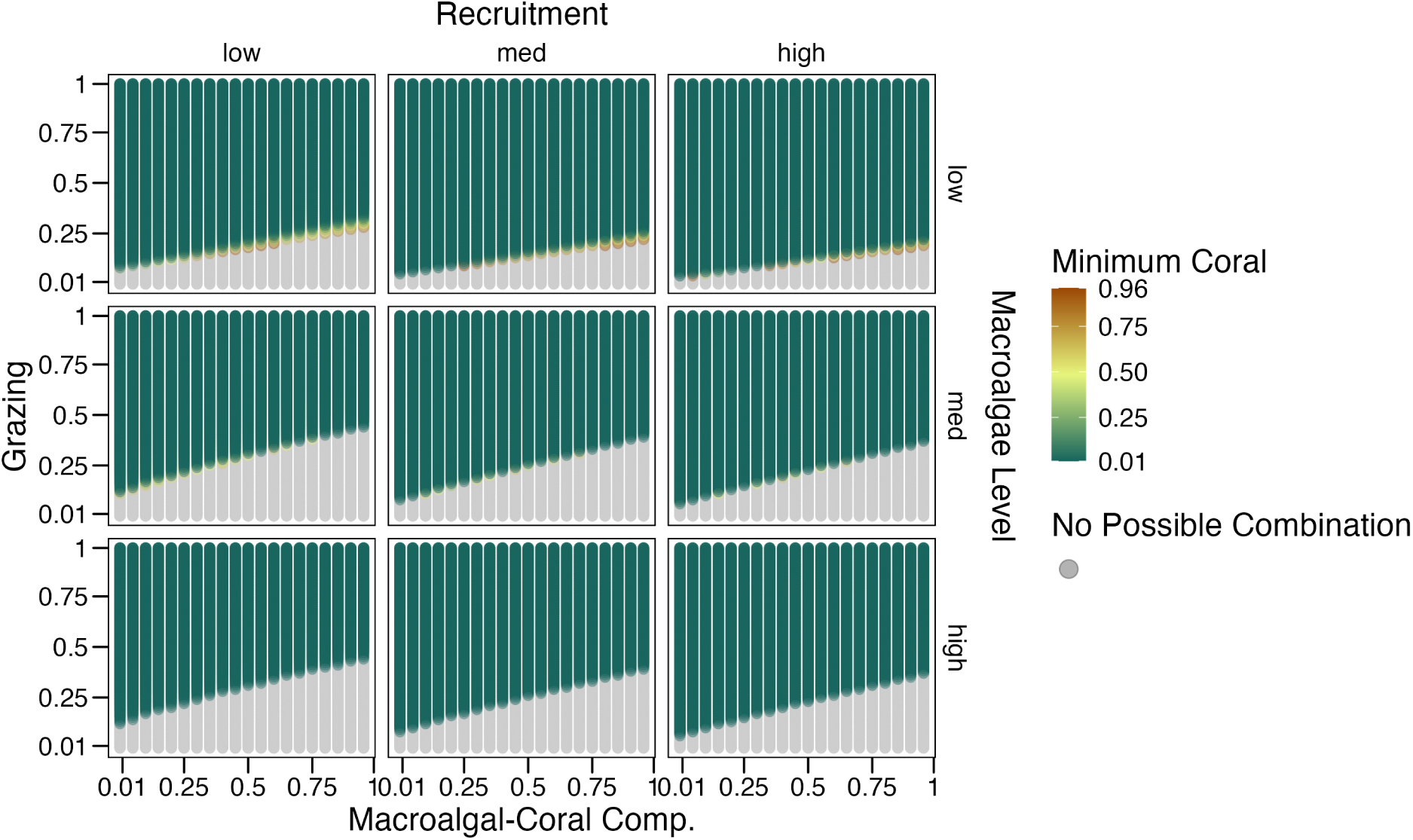
Each point represents the minimum amount of initial coral cover for the given parameter combinations, such that certain initial values of macroalgae result in the reef being in a “healthy self-sustaining coral reef state” (i.e., trending towards a stable state with *>*30% coral cover & *<*30% macroalgae cover). The values on the right y-axis depict whether the initial values of macroalgae considered are low (0-33%), medium (33-66%) or high (>66%). Light grey points indicate that no possible parameter combination or initial condition results in the reef being in a “healthy self-sustaining coral reef state”.

